# New phylogenomic analysis of the enigmatic phylum Telonemia further resolves the eukaryote tree of life

**DOI:** 10.1101/403329

**Authors:** Jürgen F. H. Strassert, Mahwash Jamy, Alexander P. Mylnikov, Denis V. Tikhonenkov, Fabien Burki

## Abstract

The broad-scale tree of eukaryotes is constantly improving, but the evolutionary origin of several major groups remains unknown. Resolving the phylogenetic position of these ‘orphan’ groups is important, especially those that originated early in evolution, because they represent missing evolutionary links between established groups. Telonemia is one such orphan taxon for which little is known. The group is composed of molecularly diverse biflagellated protists, often prevalent although not abundant in aquatic environments. Telonemia has been hypothesized to represent a deeply diverging eukaryotic phylum but no consensus exists as to where it is placed in the tree. Here, we established cultures and report the phylogenomic analyses of three new transcriptome datasets for divergent telonemid lineages. All our phylogenetic reconstructions, based on 248 genes and using site-heterogeneous mixture models, robustly resolve the evolutionary origin of Telonemia as sister to the Sar supergroup. This grouping remains well supported when as few as 60% of the genes are randomly subsampled, thus is not sensitive to the sets of genes used but requires a minimal alignment length to recover enough phylogenetic signal. Telonemia occupies a crucial position in the tree to examine the origin of Sar, one of the most lineage-rich eukaryote supergroups. We propose the moniker ‘TSAR’ to accommodate this new mega-assemblage in the phylogeny of eukaryotes.

## Introduction

The eukaryote tree of life is one of the oldest working hypotheses in biology. Over the past 15 years, this tree has been extensively reshaped based on transcriptomic and genomic data for an ever-increasing diversity of protists (i.e., microbial eukaryotes), and the development of more realistic models of evolution (Lartillot et al. 2007; Quang et al. 2008; Yabuki et al. 2014; Katz 2015; Burki et al. 2016; Brown et al. 2018; Cavalier-Smith et al. 2018; Wang et al. 2018). The current picture of the tree sees most lineages placed into a few redefined ‘supergroups’ – a taxonomic rank-less concept that was devised as inclusive clades reasonably supported by molecular and/or morphological evidence (Simpson and Roger 2002; Simpson and Roger 2004; Keeling et al. 2005). As few as eight supergroups are currently recognized, sometimes arranged in even higher-order divisions assuming the root of the tree falls outside of these mega-clades: Diaphoretickes, including Sar, Haptista, Cryptista, Archaeplastida; Amorphea, including Amoebozoa and Obazoa; Excavata, including Discoba and Metamonada (Adl et al. 2012; Burki 2014; Simpson et al. 2017). Importantly, this view of the tree remains a working hypothesis, and whilst most groups have been robustly and consistently supported, others have not and await consensual evidence (this is mainly the case for Archaeplastida and Excavata) (Burki et al. 2016; Heiss et al. 2018).

In addition to these groups, the backbone of the tree also contains many protist lineages that have long been known from early morphological description but remain without clear evolutionary affinities, as well as a few newly discovered deeply diverging lineages (Burki et al. 2009; Yabuki et al. 2014; Cavalier-Smith et al. 2015; Janouškovec et al. 2017; Heiss et al. 2018). Given this backbone, one outstanding open issue to further resolving the eukaryotic tree is to establish the branching order among all supergroups, including orphan taxa. Because many orphans are likely early diverging phyla and act as evolutionary intermediates, finding their phylogenetic position can profoundly impact our understanding of ancient eukaryote history (Brown et al. 2013; Yabuki et al. 2014; Burki et al. 2016); this was for example recently demonstrated by the establishment of ‘CRuMs’, a novel deeply branching supergroup-level clade most likely sister to Amorphea (Brown et al. 2018).

Telonemia is one such orphan taxon that has proven difficult to place in eukaryote-wide phylogenies in spite of several molecular attempts. Containing only two named species but proposed to form a phylum of its own (Shalchian-Tabrizi et al. 2006), this taxon has been shown by environmental sequencing to have well diversified in marine and freshwater environments (Shalchian-Tabrizi et al. 2007; Bråte et al. 2010). Telonemids are often detected in these aquatic environments, and although they are usually not very abundant, they are thought to play significant ecological roles as biflagellated heterotrophic predators of bacteria and small-sized eukaryotes (Klaveness et al. 2005). The ultrastructure of the group revealed specific features, such as a multilayered subcortical lamina and complex unique cytoskeleton, but also structures resembling synapomorphies in other groups, such as peripheral vacuoles similar to the cortical alveoli in Alveolata, or tripartite tubular hairs on the long flagellum similar to the mastigonemes in Stramenopila (Klaveness et al. 2005; Shalchian-Tabrizi et al. 2006). The first molecular phylogenies based on up to four genes provided only few clues as to the position of the group, other than it was not closely related to any known groups (Shalchian-Tabrizi et al. 2006). Later, more substantial analyses based on larger datasets containing 127–250 genes also failed to pinpoint with confidence the origin of Telonemia, but this was likely due at least in part to the small number of genes sampled from only one species (*Telonema subtile*) (Burki et al. 2009; Burki et al. 2012; Burki et al. 2016).

In this study, we tested whether better gene coverage and taxon sampling for Telonemia using a phylogenomic approach can help to infer with confidence the phylogenetic position of this important group of eukaryotes. To this effect, we established cultures for three telonemid species, one of which corresponded to the described *T. subtile*, and generated high-quality transcriptomes for all. The availability of different datasets from closely related species allowed us to robustly identify the correct copies from contaminant sequences, a task that is otherwise not easy with single datasets for orphan lineages. Our analyses unambiguously place Telonemia in a sister position to the supergroup Sar, forming a new mega-assemblage that further resolves the tree of eukaryotes.

## Results

### Improved dataset and taxon selection

All genes considered in this study were carefully inspected by reconstructing phylogenies to evaluate orthology and detect contamination. Sequences were removed if their positions in gene trees conflicted with an initial species tree based on an intermediate concatenated alignment (i.e., from early rounds of cleaning, see Materials and Methods); this initial species tree globally corresponded to a consensus phylogeny of eukaryotes (e.g., (Burki 2014)). For the telonemids, at least two sequences from different datasets were required to group together in gene trees in order to be retained. This requirement became possible by the addition of new telonemid datasets and greatly lowered the risk that unidentified contaminants were wrongly assigned to Telonemia. Sequences corresponding to the bodonid prey *Procryptobia sorokini*, easily identifiable in gene trees from the presence of other related taxa (e.g., *Neobodo designis, Bodo saltans*) were reassigned accordingly. Overall, 248 genes were retained for further analyses. The sequence coverage for the three telonemid OTUs ranged between 58% and 99% (Supplementary Table 1), which constitutes a large improvement over previous studies (e.g., (Burki et al. 2016)). These new telonemid OTUs were placed in the broad context of eukaryote evolution. Given the taxonomic distribution of available genome and transcriptome datasets, and the tentative placement of Telonemia in previous phylogenomic works (Burki et al. 2009; Burki et al. 2012; Burki et al. 2016), special attention was made to include representatives of all main lineages for the ‘Diaphoretickes’ part of the tree when sufficient data is available. The final taxon sampling contained 109 OTUs selected based on their taxonomic affiliation and data coverage, excluding faster-evolving lineages when the selection of shorter branches was possible (fig. 1 and Supplementary Table 1).

**Figure 1.**
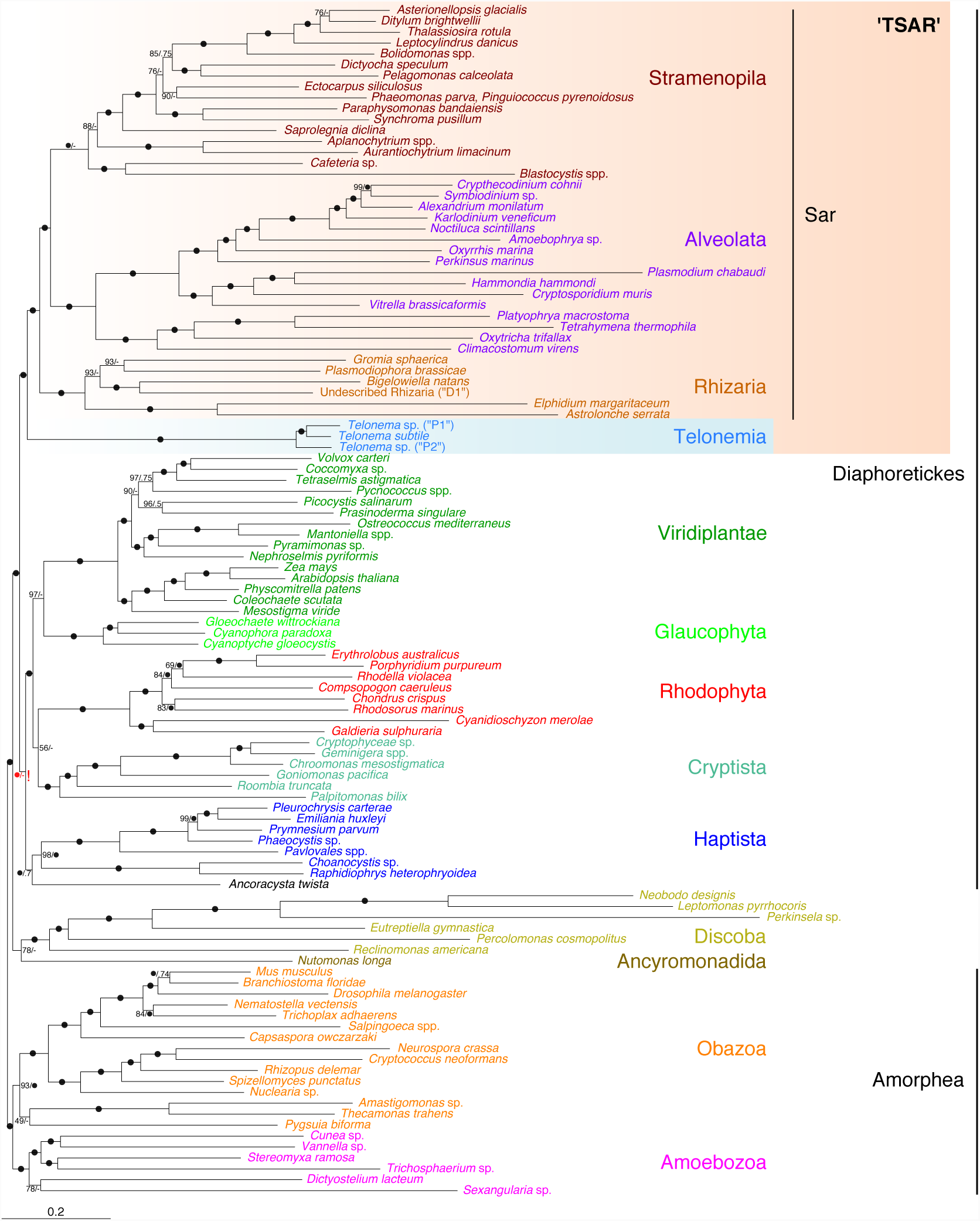
Maximum Likelihood tree showing the phylogenetic position of Telonemia within eukaryotes. The tree corresponds to the best ML tree inferred from a concatenated alignment of 248 protein-coding genes using the LG+C60+G+F-PMSF model. Support at nodes is derived from 100 bootstrap (BP) replicates and Bayesian posterior probabilities (PP). Black circles represent maximum support in both analyses. A number appended to a black circle indicates full support in only one of the two analyses (i.e., BP/PP). Nodes with dashes (-) indicate an alternative branching in the Phylobayes tree (see fig. S1 and fig. S2). The ambiguous position of Haptista with this dataset is highlighted with a red exclamation mark (see text for more information).

### Tree inference and testing

The main analyses of the 248 genes/109 OTUs supermatrix consisted in ML and Bayesian tree reconstructions using site-heterogeneous mixture models of evolution, specifically LG+C60+F+Γ4 with PMSF profiles and CAT+GTR+Γ4, respectively. These models have been repeatedly shown to be more robust against homoplasic positions, which are important to take into consideration especially along the most ancient branches of the eukaryotic tree (Lartillot and Philippe 2008; Quang et al. 2008; Wang et al. 2018). As shown in Figure 1, both models recovered a similar overall topology, with most currently accepted supergroups receiving maximal support (e.g., Sar, Cryptista, Haptista, Discoba, Obazoa, Amoebozoa). Most interestingly, in both analyses the position of Telonemia was identical and unequivocal, placed within ‘Diaphoretickes’ as sister to the Sar group (BP = 100%, PP = 1.0). Cryptista was robustly inferred in a clade including the members of Archaeplastida, but the branching pattern in this clade was unclear: whilst the LG+C60+F+Γ4+PMSF tree remained unresolved (BP = 56% for the grouping of Cryptista with Rhodophyta), the CAT+GTR+Γ4 model recovered a monophyletic Archaeplastida with 1.0 PP (fig. 1). The position of Haptista (and its possible sister taxa, *Ancoracysta twista*; see (Janouškovec et al. 2017)) also remained ambiguous, showing a sister relationship to the Archaeplastida + Cryptista clade with the LG+C60+F+Γ4+PMSF model (BP = 100%) but corresponding to an unconverged node in the CAT+GTR+Γ4 analysis (two chains supporting the named relationship, and two chains supporting a sister relationship to Telonemia + Sar; for details, see fig. S1 and fig. S2).

Therefore, the relationships within ‘Diaphoretickes’ show two strongly supported blocks, Telonemia + Sar and Cryptista + Archaeplastida (whose monophyly remains ambiguous), as well as a third group, Haptista, whose position is unclear (fig. 1). In order to test for the consistency of these results, we performed two alterations of the main 248 genes/109 OTUs supermatrix. First, we evaluated the persistence of the phylogenetic signal for the Telonemia + Sar and Cryptista + Archaeplastida groupings when progressively decreasing the number of genes used to infer the trees (fig. 2A). Random subsamples of the total 248 genes were generated without replacement in 20% decrements (80% to 20%), with a replication scheme that ensured a 95% probability of sampling every gene in each subset (Brown et al. 2018). Each replicate was then subjected to ML tree reconstruction under the LG+C60+F+Γ4 model and the group support monitored with ultrafast bootstrap approximation (Hoang et al. 2018). Both groups (i.e., Telonemia + Sar and Cryptista + Archaeplastida) showed remarkable stability until as few as 60% of the genes were subsampled, corresponding to an alignment length of 149 genes (in average 34,588 AA positions), but started to crumble with fewer genes (fig. 2A). None of the other tested groups were as robust against the decreasing number of genes, with the exception of Sar (used as control) that remained maximally supported throughout the experiment (fig. 2A).

**Figure 2.**
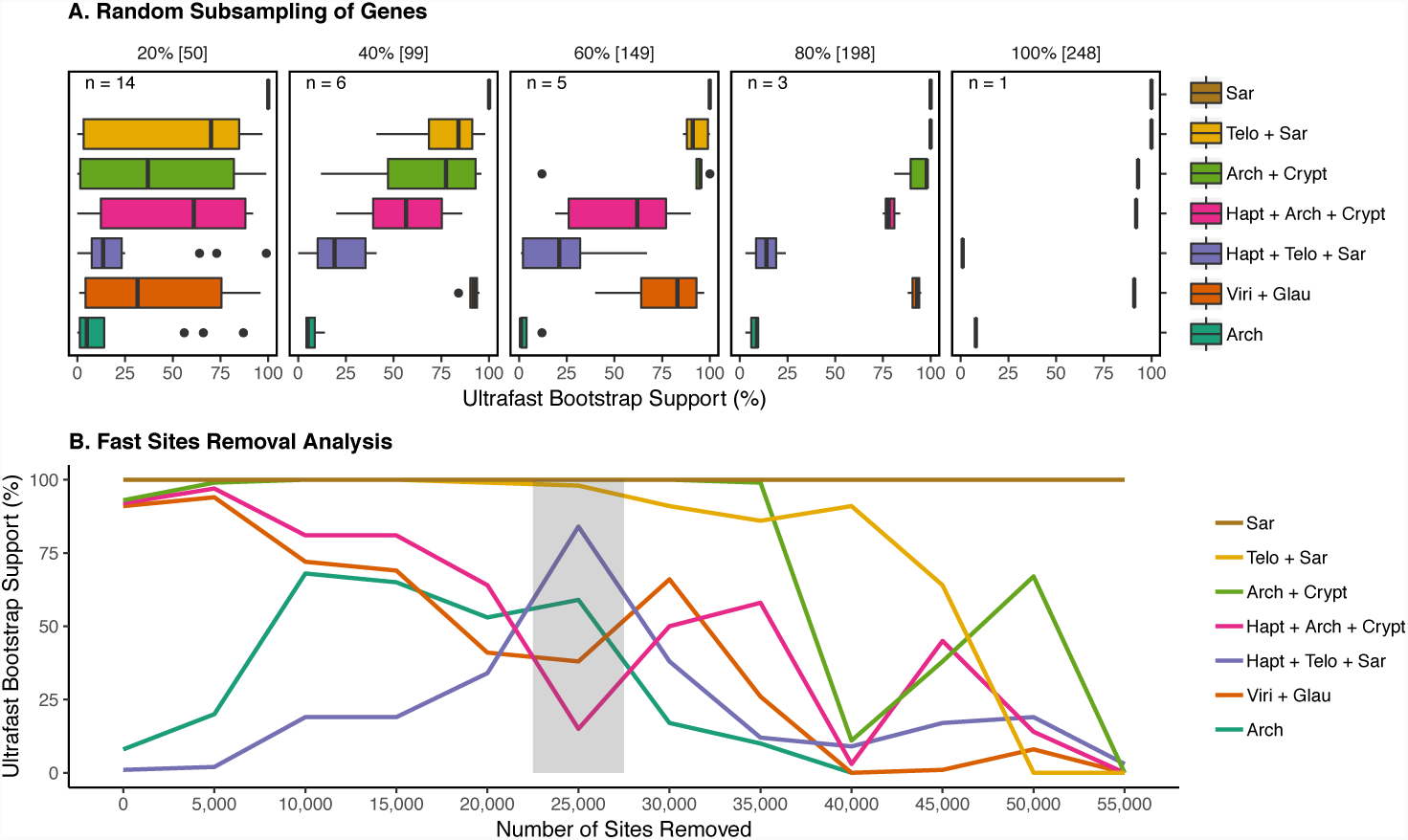
Gene subsampling and fast-evolving sites removal analyses. (A) Ultrafast bootstrap approximation (UFBoot) for selected nodes of interest using the full supermatrix (100%) and subsets of randomly sampled genes (20% to 80%; see Materials and Methods). The variability of support values is shown by Box-and-Whisker plots. (B) Ultrafast bootstrap approximation (UFBoot) for the same nodes as in (A) using the full supermatrix (0 sites removed) and subsets from which fast-evolving sites were removed in 5,000 increments (see Materials and Methods). The grey box indicates the reduced dataset which recovered a different position for Haptista. Arch – Archaeplastida, Crypt – Cryptista, Glau – Glaucophyta, Hapt – Haptista, Telo – Telonemia, Viri – Viridiplantae.

The second alteration was the removal of the fastest evolving sites from the full supermatrix, which are more likely to be homoplasic and thus leading to systematic errors due to model misspecification (Rodriguez-Ezpeleta, Brinkmann, Roure, et al. 2007). Sites were sorted from fastest to slowest and progressively removed in 5,000 sites increments until only 3,469 sites were left. Each sub-dataset was again used in ML tree reconstruction (LG+C60+F+Γ4 model) with ultrafast bootstrap approximation (Hoang et al. 2018). The Telonemia + Sar relationship remained strongly supported until 25,000 sites were removed, after which the support was progressively lost but not replaced by an alternative supported relationship (fig. 2B). Similarly, the Cryptista + Archaeplastida group retained strong support until the removal of 35,000 sites with no other supported grouping emerging when more sites were removed. In contrast, the position of Haptista was markedly altered by the removal of fast-evolving sites. Starting from a maximally supported sister relationship to the Cryptista + Archaeplastida group (fig. 1 and fig. 2B), Haptista shifted to a sister position to the Telonemia + Sar group after the removal of 25,000 sites (82% UFBoot), before reverting again to its original position with further site removal albeit with no real statistical significance (fig. 2B). In order to further assess the strength of the Telonemia + Sar + Haptista relationship, we ran a Bayesian analysis with the CAT+F81 model (Lartillot et al. 2013) using the 25,000 sites-removed dataset. Interestingly, the two converged chains were fully consistent with the ML tree and the position of Haptista as sister to Telonemia + Sar received maximal PP (fig. 3 and fig. S3).

**Figure 3.**
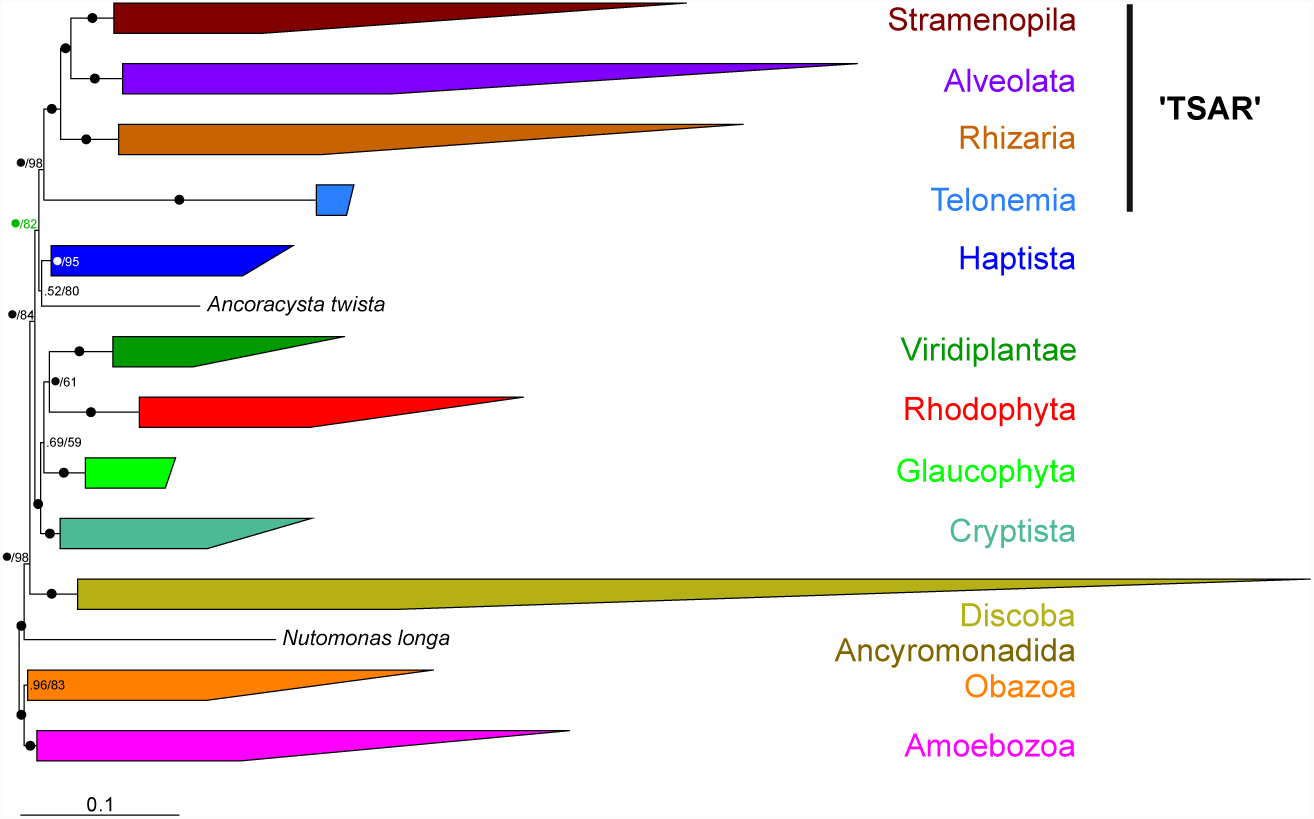
Simplified Phylobayes consensus tree inferred from a reduced concatenated alignment (25,000 fast-evolving sites removed) of the 248 protein-coding genes under the CAT+F81+Γ4 model (for a full version showing all taxa, see fig. S3). Node support is given by Bayesian posterior probabilities (PP) and ultrafast bootstrap approximation (UFBoot) inferred from a ML analysis (LG+C60+G+F model) using the same sub-alignment (i.e., 25,000 sites removed; see text and fig. 2). Nodes with black circles are fully supported in both analyses. A number appended to a black circles indicates full support in only one of the two analyses (i.e., PP/UFBoot). Note the complete congruence between the Bayesian and ML analyses (all nodes are recovered by both methods), including the position of Haptista (labelled in green).

## Discussion

The position of Telonemia in the tree of eukaryotes has been debated ever since the group was first described (Vørs 1992; klaveness et al. 2005; Shalchian-Tabrizi et al. 2006). The difficulty in placing Telonemia is due to a combination of several factors, including the ancient origin of the group, the sparse sampling of molecular data for group members, and the lack of closely related outgroup lineages. Earlier phylogenomic studies often placed Telonemia in the vicinity of Sar or close to cryptist or haptist lineages, but never convincingly (Burki et al. 2009; Burki et al. 2012; Katz and Grant 2015; Burki et al. 2016). Here, we tested whether improving the gene coverage in our phylogenomic supermatrix and adding more telonemid representatives help to infer the origin of this group with more confidence. The results of the phylogenetic analyses show that this is the case: for the first time, the placement of Telonemia in the global phylogeny of eukaryotes receives strong and consistent support across multiple analyses (ML and Bayesian analyses using site-heterogeneous models, fig. 1). Telonemia was inferred as the most closely related lineage to the Sar supergroup. This position is robust against variations in the gene sets although it requires a substantial amount of data to be recovered (typically above 150 genes), and is stable when the most saturated sites are removed so it is unlikely to be due to convergence from undetected multiple substitutions (fig. 2). To acknowledge the strength of this relationship and facilitate further discussions, we introduce the constructed name ‘TSAR’ to refer to the Telonemia + Sar grouping. We propose that TSAR forms a provisional supergroup of eukaryotes.

The hypothesis of TSAR has multiple implications. Most importantly, it confirms that Telonemia is a *bona fide* eukaryotic phylum, at the same phylogenetic level as other major phyla such as the Sar members (Stramenopila, Alveolata, and Rhizaria) (Burki et al. 2007; Hackett et al. 2007; Rodriguez-Ezpeleta, Brinkmann, Burger, et al. 2007). A unique combination of ultrastructural cellular characteristics, such as a complex cytoskeleton, peripheral vacuoles and tubular tripartite hairs, suggested that Telonemia might be distinct from all better studied major eukaryotic groups but possibly evolutionary connected to lineages such as Alveolata or Stramenopila (Shalchian-Tabrizi et al. 2006). Here, we show that Telonemia indeed shares a most recent common ancestry with Sar. Consequently, it is possible that Telonemia reflects the ancestral state of the Sar group before the cytoskeleton was reduced and homologous ultrastructural features differentially lost in derived lineages (namely the loss of tripartite tubular flagellar hairs in Alveolata, alveoli in Stramenopila, and both structures in Rhizaria). Telonemia thus appears crucial for understanding the early evolution of the Sar group, a globally distributed and highly diverse assemblage of eukaryotes likely containing the majority of protist lineages (del Campo et al. 2014; de Vargas et al. 2015; Grattepanche et al. 2018).

In addition to TSAR, our analyses strongly recovered another major clade in Diaphoretickes, including Archaeplastida members (not consistently monophyletic) and Cryptista (AC clade; fig. 1). This association has been recovered before in taxon-rich phylogenomics, and its implication for eukaryote evolution discussed (Burki et al. 2016). However, the AC clade still awaits confirmation because other recent studies displayed alternative relationships, albeit with no conclusive support and poorer taxon sampling for the relevant groups (Yabuki et al. 2014; Cavalier-Smith et al. 2015; Katz and Grant 2015; Cavalier-Smith et al. 2018). Our analyses also failed to consistently place Haptista in the tree. Based on the full dataset, ML and some unconverged Bayesian chains favoured a close relationship to the AC clade (fig. 1). But after the removal of the 25,000 fastest-evolving sites, both ML and converged Bayesian analyses supported the placement of Haptista and TSAR together (fig. 3). The position of Haptista has varied dramatically with conflicting confidence support (e.g., (Yabuki et al. 2014; Katz and Grant 2015; Burki et al. 2016; Cavalier-Smith et al. 2018)). Hence, moving forward to improve the deep structure of Diaphoretickes, it will be crucial to specifically address the causes for the lack of consensus for the position of Cryptista, Haptista, and the monophyly of Archaeplastida.

Despite these lasting uncertainties surrounding the relationships within Diaphoretickes, we present here consistent results that bring us one step closer to a fully resolved eukaryotic tree of life. Telonemia was once a notorious rogue lineage, but using new transcriptome data for this important but neglected phylum, we show that it now occupies a robust position in the tree as part of an expanded supergroup (TSAR). We thus advise that Telonemia should be routinely included in future works addressing the broad-scale evolution of eukaryotes, in the same way as other major lineages are included. More generally, this study follows a series of recent phylogenomic investigations that successfully placed in the tree several orphan taxa (Brown et al. 2013; Yabuki et al. 2014; Burki et al. 2016; Janouškovec et al. 2017; Brown et al. 2018). This demonstrates that combining adequate datasets and phylogenomics can go a long way towards filling important evolutionary gaps. With the maturation of culture-free genomic solutions that allow to efficiently work with very little starting material, we expect that most long-standing mystery taxa will soon fill the tree of life and contribute to a better understanding of ancient eukaryote evolution.

## Materials and Methods

### RNA Extraction, Sequencing, and Assembling

Clonal cultures of three telonemid species were isolated from Arctic surface waters of the Kara Sea, N 75.888, E 89.508, total depth: 52 m, water temperature: 2.3 °C. Isolates were propagated on the bodonid *Procryptobia sorokini* Frolov, Karpov, and Mylnikov, 2001 strain B-69 grown in Schmalz-Pratt’s medium by using the bacterium *Pseudomonas fluorescens* as food (Tikhonenkov et al. 2014).

For cDNA preparation, cells were harvested following peak abundance after eating most of the prey. Cells were collected by centrifugation (2,000 × g, room temperature) on the 0.8 *μ*m membrane of Vivaclear Mini columns (Sartorium Stedim Biotech Gmng, Germany, Cat. No. VK01P042). Total RNA was extracted using the RNAqueous®-Micro Kit (Invitrogen™, Cat. No. AM1931) and was converted into cDNA prior to sequencing using the SMART-Seq® v4 Ultra® Low Input RNA Kit (Takara Bio USA, Inc., Cat. No. 634888). Sequencing was performed on the Illumina MiSeq platform with read lengths of 300 bp using the NexteraXT protocol (Illumina, Inc., Cat. No. FC-131-1024) to construct paired-end libraries.

Reads were trimmed with Trimmomatic v. 0.36 (Bolger et al. 2014) using the following parameters: LEADING:5, TRAILING:5, SLIDINGWINDOW:5:16, MINLEN:80. Potential vector sequences were detected (and if applicable removed) by BLAST searches against the UniVec database (e-value: 1e-4). Cleaned reads were assembled and translated into amino acids with Trinity v. 2.4.0 and TransDecoder, respectively (default settings (Haas et al. 2013)). In addition to the reads obtained in our study, publicly available reads of other taxa were trimmed, assembled, and translated as described above but with slightly different settings for the trimming procedure (LEADING:3, TRAILING:3, SLIDINGWINDOW:4:15, MINLEN:36).

### Phylogenomic Dataset Construction

A previously published dataset comprising 263 genes and 234 taxa (Burki et al. 2016) was used as starting point for this study. New taxa (sources: ensemblgenomes.org, imicrobe.us/#/projects/104, ncbi.nlm.nih.gov, onekp.com) were added by the following successive steps: 1) Protein sequences of each taxon were clustered with CD-HIT (Fu et al. 2012) using an identity threshold of 85%; 2) Homologous copies were retrieved by BLASTP searches using the 263 genes as queries (e-value: 1e-20; coverage cutoff: 0.5); 3) Phylogenetic trees were constructed and carefully inspected in order to detect and remove contaminants and select orthologous copies. An automated colouring scheme for the trees was developed for easier visual inspection in FigTree v. 1.4.3 (http://tree.bio.ed.ac.uk/software/figtree/). Three rounds of inspection were performed, first to detect deep paralogs and progressively refined to remove contamination and select orthologs. Sequences were aligned with MAFFT v. 7.310 (Katoh and Standley 2013) using the -auto option (first round) and with MAFFT L-INS-i using the default settings (second and third round), and filtered with trimAL v. 1.4 (Capella-Gutiérrez et al. 2009) using a gap threshold of 0.8 (all three rounds). FastTree v. 2.1.10 (Price et al. 2010) employing -lg -gamma plus options for more accurate performances (first round) and RAxML v. 8.2.10 (Stamatakis 2014) employing PROTGAMMALGF and 100 rapid bootstrap searches (second and third round) were used to infer single gene Maximum Likelihood (ML) trees. Three genes (FTSJ1 – RNA methyltransferase 1, tubb – tubulin beta, and eif5A – eukaryotic translation initiation factor 5A) were excluded from the base dataset for further analyses due to suspicious clustering of major groups; e.g., in FTSJ1, a duplication of the entire Sar group could be detected, and in tubb, an ambiguous topology was observed due to duplications within the Stramenopila and Opisthokonta. Twelve other genes were removed because they lacked telonemid sequences.

The final dataset comprised 248 genes and 293 taxa. An unaligned version of these genes was then subjected to PREQUAL v. 1.01 (Whelan et al. 2018), a new software to remove or mask stretches of non-homologous characters in individual sequences. A posterior probability threshold of 0.95 (-filterthresh) was used. MAFFT G-INS-i with the VSM option (--unalignlevel 0.6) was used for alignment, and ambiguously aligned sites were filtered out with BMGE v. 1.12 (Criscuolo and Gribaldo 2010) with the parameters -g 0.2, -b 5, -m, and BLOSUM75. Partial sequences belonging to the same taxon that did not show evidence for paralogy or contamination on the gene trees were merged. Of the initial 293 taxa, a reduced set of 148 taxa was selected to allow computationally intensive analyses based on i) knowledge of sequence divergence (slower-evolving species were retained when possible), ii) to represent all major eukaryotic groups with sufficient data (amino acid positions in supermatrix >55%); of all major eukaryotic lineages with genomic or transcriptomic data, only Picozoa was deemed of insufficient coverage, and iii) sequence completeness among the genes. Furthermore, in order to reduce missing data, several strains and species complexes were combined when monophyly inferred from a preliminary concatenated alignment was unambiguous (tree not shown but see Supplementary Table 1 for a list of 109 taxa and chimera that were used as operational taxonomic units; OTUs). Finally, all 248 genes were concatenated into a supermatrix (58,469 amino acid positions) with SCaFos v. 1.25 (Roure et al. 2007).

### Phylogenomic Analyses

ML trees were inferred from the supermatrix using IQ-TREE v. 1.6.3 (Nguyen et al. 2015). Following the Akaike Information Criterion (AIC) and the Bayesian information criterion (BIC), the best fitting model was the LG+C60 mixture model with stationary amino acid frequencies optimized from the dataset (F) and four discrete gamma (Γ) categories for rate heterogeneity (LG+C60+F+Γ4). The tested models were site-homogeneous LG +/- F, I, Γ4; site-heterogeneous LG +/- F, Γ4 and C10–C60. The best ML tree for the full dataset was used as guide tree under the same model to estimate the posterior mean site frequencies (PMSF; (Wang et al. 2018)) model, which was used for further tree reconstructions (LG+C60+F+Γ4+PMSF). Support values for the best LG+C60+F+Γ4+PMSF tree were obtained using non-parametric bootstrap (BP) with 100 replicates.

Bayesian analyses were inferred using PHYLOBAYES-MPI v. 1.8 (Lartillot et al. 2013) with the CAT+GTR+Γ4 model. Constant sites were removed to decrease running time (-dc). Four independent Markov Chain Monte Carlo (MCMC) chains were run for >6,300 generations (all sampled). For each chain, the burnin period was chosen after monitoring the evolution of the log-likelihood (Lnl) at each sampled point, removing the generations before stabilization (generally 1,250) prior to computing a 50% majority-rule consensus tree of all chains using *bpcomp*. Global convergence between chains was assessed by the maxdiff statistics measuring the discrepancy in posterior probabilities (PP). Unfortunately, global convergence was never achieved in any combination of two chains; this is a known issue of PHYLOBAYES, which is especially acute with taxon-rich datasets and has been reported on many occasions (e.g., (Katz and Grant 2015; Burki et al. 2016; Kang et al. 2017)). The discrepancies between chains concerning nodes under scrutiny in this study were labelled as unresolved.

### Gene Subsampling and Fast-evolving Sites Removal

From the 248 gene dataset, random subsampling without replacement of 80% to 20% (in 20% increments) of genes were performed with replications (3, 5, 6, 14 replicates in each subset, respectively); this allowed a >95% probability that every gene is represented at least once (following an approach described in Brown et al. 2018). The genes in each subset were then concatenated with SCaFoS. Fast-evolving sites were estimated with IQ-TREE v. 1.6.5 (option -wsr) giving the PMSF tree as fixed tree topology and using the LG+C60+F+Γ4 model. Sites were then removed in 5,000 increments using a Perl script. For all the subsamples with reduced number of genes or sites, phylogenetic trees with ultrafast bootstraps (1,000 replicates) were inferred using IQ-TREE (LG+C60+F+Γ4) with the -bb, wbtl, and -bnni flags employed. One sub-alignment (25,000 sites removed) was used in a Bayesian tree inference using the site-heterogeneous CAT+F81+Γ4 model as implemented in PHYLOBAYES-MPI v. 1.8 (Lartillot et al. 2013). We did not use here the CAT+GTR+Γ4 model for computational tractability. Two independent MCMC chains were run for >6,300 cycles. Before computing the majority-rule consensus tree from the two chains combined, the initial 2,500 trees in each MCMC run were discarded as burnin after checking for convergence in likelihood and in clade posterior probabilities (maxdiff < 0.3).

## Acknowledgments

This work was supported by a grant from Science for Life Laboratory available to FB, which covered salaries of JFHS and MJ, and experimental expenses. The work of DVT and APM was supported by the Russian Science Foundation (no. 18-14-00239). DVT thanks Dr. Natalia Kosolapova (IBIW RAS) for help with sample collection. Computational resources were provided in part by the Swedish National Infrastructure for Computing through UPPMAX under project SNIC 2018/8-192.

## Data deposition

All new transcriptomic data have been submitted to the SRA database under accession number SRP150904 (https://www.ncbi.nlm.nih.gov/sra/SRP150904). All single gene alignments, gene trees, and the supermatrix are available online in supplementary file TSAR_Sequence_data_and_trees.zip.

